# Neurodevelopmental clustering of gene expression identifies lipid metabolism genes associated with neuroprotection and neurodegeneration

**DOI:** 10.1101/2021.09.02.458277

**Authors:** Akiva A. Kohane, Tim R. Wood

**Author notes:** Correspondence should be addressed to A.A.K.

## Abstract

APOE variants present the strongest association with sporadic Alzheimer’s Disease. APOE is also highly expressed during neurodevelopment in the Central Nervous System (CNS) and has been shown to be neuroprotective in infancy and gestation. We explored other lipid metabolism genes to determine whether they show a similar neurodevelopmental expression trajectory and are associated with neurodegeneration. APOE was by far the most highly expressed of the lipid metabolism genes in the CNS. Two other genes, Apolipoprotein C1 (APOC1) Glutamate-Ammonia Ligase (GLUL—also known as glutamine synthetase), co-clustered with Apolipoprotein E (APOE) in its developmental trajectory in late gestation through early childhood. These three genes highlight brain structures and developmental time-windows distinct from other lipid metabolism genes. In the CNS they are primarily expressed in astrocytes and are implicated in neuroprotection and neurodegeneration.

## Introduction

APOE is a lipoprotein involved in lipid metabolism with specific variants implicated through epidemiological and in vitro evidence with neurodegeneration [1]. The variant APOE4 has been repeatedly identified as the highest risk genetic factor for sporadic Alzheimer’s disease as well as contributing to cardiovascular disease. Although primarily produced in the liver, APOE is also expressed in the CNS where it is mostly produced by astrocytes, much less by oligodendrocytres and microglia and even in neurons in specific pathological states[2]. In the CNS, APOE is the major apolipoprotein carrier of cholesterol and because of its role in generating HDL-like particles in the CNS it is also a major determinant of the distribution of phospholipids in the CNS. LDL Cholesterol itself is synthesized in the CNS from acetate and not imported from the periphery and the blood brain barrier prevents most lipids including cholesterol to enter the CNS. Lipids overall form 50% of the dry weight of the brain, second only to adipose tissue. Of that dry weight glycerophosphatides account for approximately 20% in gray matter and white matter and 78% in myelin. Cholesterol accounts for 8% in the gray matter, 15% in white matter and 20% in myelin. [3].

Given the centrality of lipids and cholesterol in the composition of the human brain, the abundance of APOE in the CNS, the growing evidence that cholesterol and lipid metabolism disorders have been implicated in several neurodevelopmental disorders some rare [4] and others more common (e.g. ADHD)[5,6], we sought to investigate which lipid metabolism genes had similar neurodevelopmental trajectories to APOE to determine which could be implicated in both neurodevelopmental processes and neurodegenerative diseases. This guilt-by-association strategy was enabled by the availability of transcriptome-wide data sets, specifically the Brainspan data set generated as part of the Allen Brain Atlas [7]. This data set provides transcriptomic measures at multiple developmental stages and in multiple defined structures of the CNS. We used an unsupervised learning/clustering approach across these tissues and developmental states to find the closest neighbors in tissue and developmental state trajectories.

## Results

### First subheading has no extra space before it

APOE is the most highly expressed gene of all the 103 gene lipid metabolism panel. As illustrated by the heatmap in Figure 1, it is expressed at one to two orders of magnitude higher levels than most of the other panel genes during embryogenesis to age 40 years. The higher levels of expression start at approximately 25 post-conceptual weeks (pcw) and continue to rise through age 2 years and then return to much lower expression levels before age 30 years as illustrated in Figure 2. Most of the neocortex follows this expression pattern but the highest APOE expression levels occur in the thalamus. The trajectory for hippocampus and striatum APOE expression follows a similar pattern but at lower levels.

**Figure 1:**
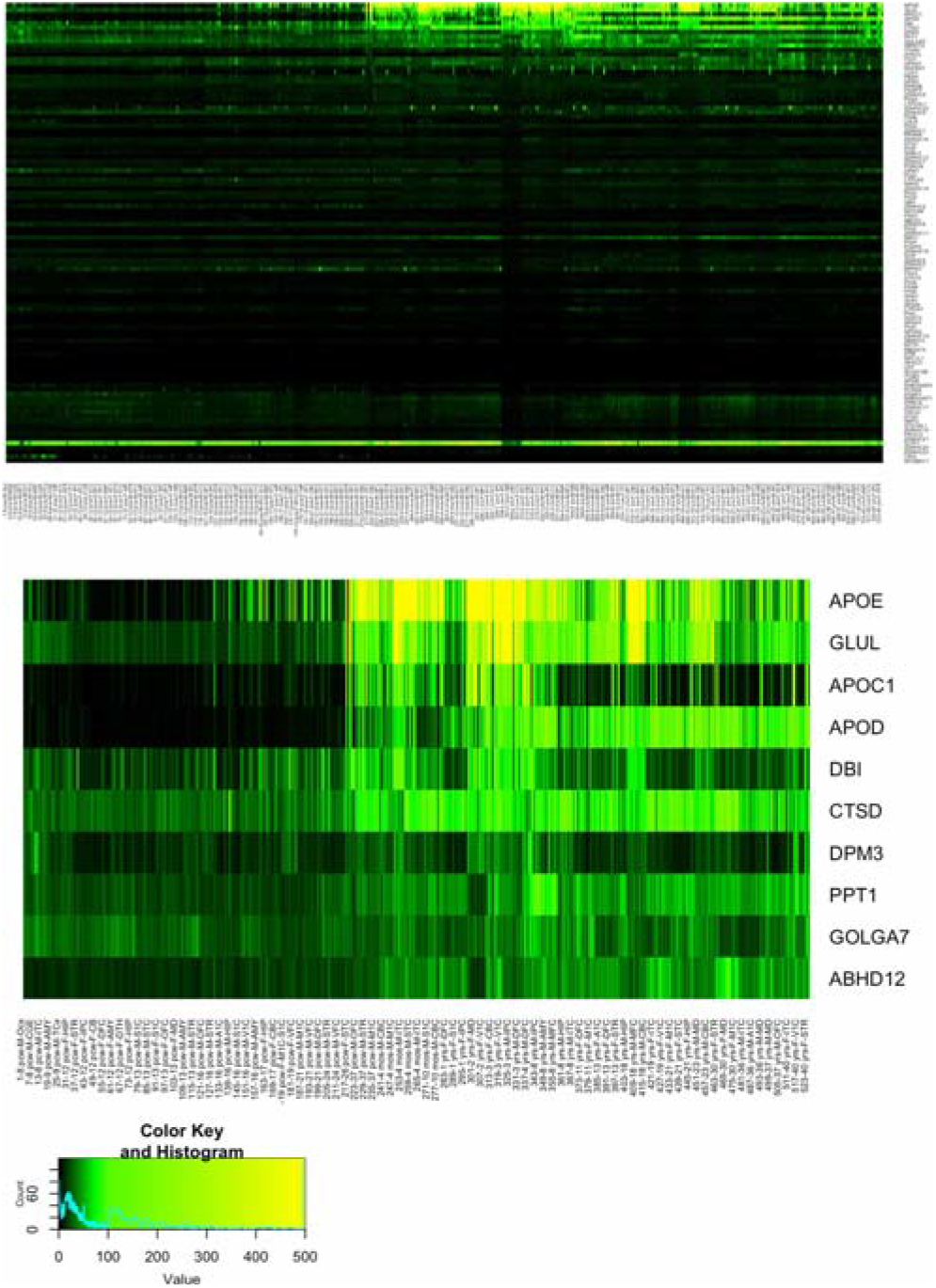
Heatmap of gene expression across lipid panel. Rows correspond to genes ordered by their coefficient in Column 0 of the NMF W matrix (descending from top to bottom) and the columns correspond to samples ordered by increasing age (left to right). The middle panel shows the top 10 genes in the topmost panel for clarity. The bottom panel is the color key and histogram of gene expression values.

**Figure 2:**
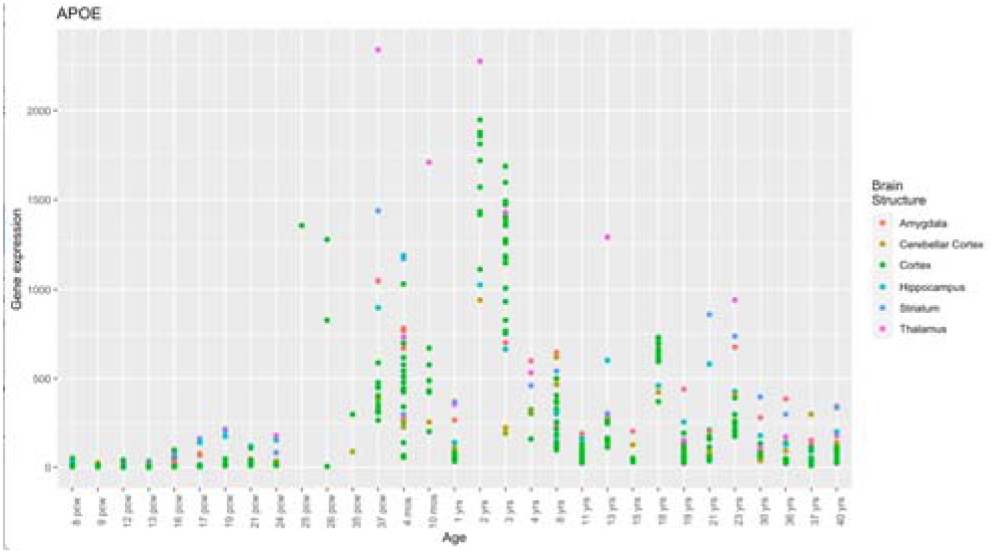
Structure- and Age-specific APOE expression.

The gene expression profiles of the lipid panel were analyzed by non-negative matrix factorization to generate two unsupervised clusters (see Methods, below). Each column in the W matrix corresponds to a cluster and each row the coefficient of a gene. APOE has the highest coefficient in Column 0 (see Supplementary Figure S1b)— cluster C0. The coefficient of APOE in C0 was two orders of magnitude higher than most of the lipid panel genes. The next most highly ranked genes in C0 are GLUL and APOC1. Like APOE their coefficients in column 0 of the W matrix Higher than in column 1. As shown in Supplementary Figure S3(a, b) they have similar trajectories to APOE. The C0 coefficient of the next highest ranked gene, APOD, is lower than that in C1 unlike the higher ranked genes (APOE, GLUL, APOC1). Therefore, it is not surprising that the APOD trajectory, across all samples, not just in C0, in Supplementary Figure S3© is markedly different from the first three trajectories described above. This is reflected in the correlation heatmap (Supplementary Figure S4) where the GLUL and APOC1 are the most tightly correlated with APOE (with **z** scores of 23.2 and 24.8 respectively) with APOD (**z** score 18.1) next most correlated.

The plots described above include all the samples in the data set. If instead we stratify the samples by their membership in C0 or C1, an obviously and consistently decreased APOE expression is seen (Supplementary Figure S2(a)) across C1 samples relative to C0 samples. This difference is also true for GLUL and APOC1 (not shown) and is associated with a highly overlapping set of samples as illustrated for APOE and GLUL in Supplementary Figure S2(b, c). A t test comparing the male to the female samples’ expression levels was not significant with p > 0.05.

Querying the GWAS catalog[13] for the associations for Alzheimer’s Disease/neurodegeneration (AD/ND) reveals the well-known association for APOE and the association for APOC1 which also has been previously reported and may also contributes to the APOE4 risk independent of its proximity to APOE [14]). There are no known significant GWAS associations between GLUL and AD/ND.

## Discussion

Given the importance of lipids as constituents of brain structure and lipid metabolism in neurodevelopment and neurodegeneration, we undertook a unsupervised clustering study of lipid metabolism genes in the developing CNS. Within the two clusters obtained across developmental time and across tissues, three genes, APOE, APOC1 and GLUL were top ranked in one cluster and with the greatest difference with respect to the second cluster. Furthermore, APOE was most tightly correlated across the entire data set with the same two genes, all with p values < 2.2 × 10^−16^.

GLUL catalyzes the production of glutamine from glutamate and ammonia. Glutamine is a central metabolite in cellular energy balance including fatty acid synthesis particularly during hypoxia [15]. In the brain GLUL, mostly produced by astrocytes, eliminates neurotoxic glutamate and neurotoxic ammonia by synthesizing glutamine which is then taken up by glutamatergic neurons. Those neurons subsequently deamidate the glutamine to generate glutamate to serve energy metabolism or as neurotransmitter.

GLUL has been shown to be neuroprotective [16] during neurodevelopment in mopping up glutamate and ammonia both of which are neurotoxic. Independently, glutamine protects against cholesterol-dependent cytolysis [17]. Further glutamine blockade increases inflammatory signatures in microglia [18]. APOE is tightly regulated by processes of nerve degeneration and regeneration [19]. AD APOE risk alleles have been associated with differential pediatric gray matter maturation [20] as have responses to insults such as traumatic injury [21,22] and infections [23]. APOC1 suppresses glial inflammatory signaling and its expression is modulated downwards by the APOE ε4 AD risk allele [14].

Regarding neurodegeneration, APOE variants have been the most studied and demonstrated to contribute more to sporadic AD than any other variants. APOC1 has also been shown through eQTL analysis to have an independent contribution to AD risk, possibly through modulating APOE expression. GLUL variants have not been associated with AD. However, a recent mendelian randomization study supports the hypothesis of a neuroprotective effect of circulating glutamine regarding AD [24]. This suggests that prior findings of elevated expression of GLUL in brains of AD patients may be a compensatory rather than a pathogenetic mechanism [25].

In summary, three genes: —APOE APOC1 and GLUL—were highly expressed and highly correlated in a study of developmental gene expression in the brain. All three rise from 25 weeks pcw to age 3, most markedly in the neocortex and thalamus. Additionally, all three are known to be expressed preferentially in astrocytes compared to other CNS cells.

All three of these genes have been shown to have neuroprotective effects during neurodevelopment. Also, all three are also associated with neurodegeneration suggesting that the neuroprotective effects in gestation may be mechanistically related to neuroprotective effects during neurodegeneration. Although this study focused on lipid metabolism, it may be that other neurodevelopmental neuroprotective genes are also important in neurodegenerative processes.

## Limitations

The heuristics developed to identify the relevant brain-expressed, developmental-disorder associated genes are described in Methods (below). Other genes might be identified by different heuristics and may also share the APOE, GLUL, APOC1 group trajectories. Further, there may be other characteristics of the patients that were not recorded in the metadata that might explain the observed clustering in the PCA and subsequently analyzed by NMD.

## Author contributions

A.A.K. conducted all the analyses and wrote the manuscript. T.R.W. suggested analytic techniques to cluster the expression data. He also provided insight regarding the connection between developmental signatures and neuroprotection vs neurodegeneration.

## Methods

### Lipid Panel Selection

We curated a list of 103 genes—the “lipid panel”—(see Supplementary Table 1) involved in lipid metabolism which met the following criteria: 1) They were expressed in the brain as reported by a tissue-wide transcriptomic survey [8] with an expression level of at least 1 RPKM 2) They were associated in PubMed with lipid metabolism (using the Chilibot tool [9]) 3) They were associated with neurodevelopmental disorders as reported by the Online Mendelian Inheritance in Man database [10]

### Gene Expression Data

Expression levels of each of those genes across multiple brain structures across development as measured in the Brainspan developmental transcriptome project (https://www.brainspan.org/) were downloaded. These gene-specific expression measures were previously described [7]. Some brain structures were sparsely represented across the Brainspan database and therefore therefore they were aggregated according to the mapping below.

### Analysis

The gene expression measures were unit standard deviation normalized (see the Github repository at bit.ly/akivalipo for a Jupyter notebook with these analyses). Principal component analyses (PCA) revealed two clusters separated across PC1 (Supplementary Figure 1a). Non-Negative Matrix Factorization (NMF) [11] was used to determine the contributions of each gene to the clustering of the samples. Specifically, the original expression matrix **V** (with the columns representing samples and rows genes) is approximated with two smaller matrices **W** and **H** such that:

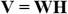

The W matrix can be viewed as an archetype for each sample cluster with each column containing the coefficients for each gene for that cluster. NMF as implemented by the Python sci-kit library [12] was run with k=2 as informed by the two clusters seen in the PCA. Therefore, in this instance there are two columns in the W matrix. These are shown (Supplementary Figure 1b) for the top 10 genes ranked by their coefficients in column 0, along with the corresponding coefficients in column 1 and the absolute difference between the two columns shown in column 3.

**Supplementary Figure S1:**
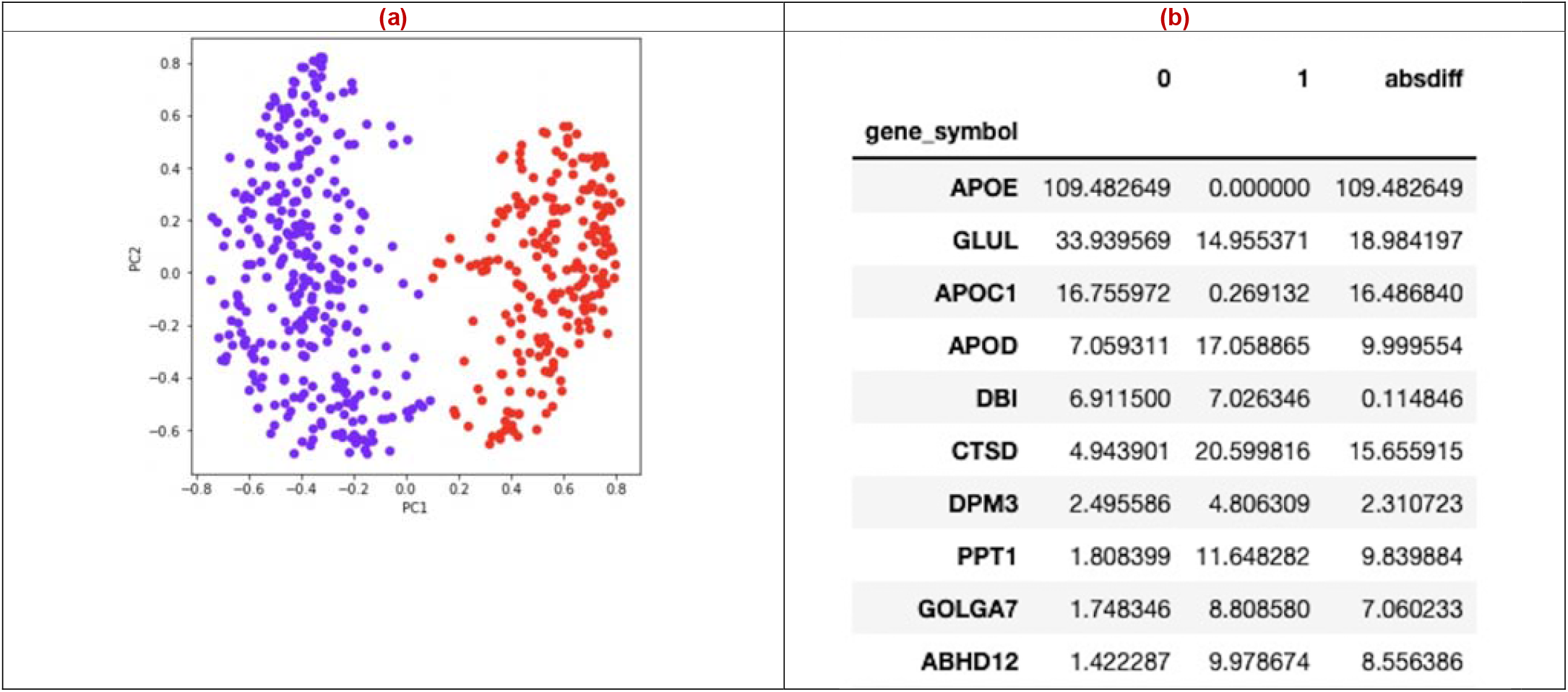
Clustering of samples by expression values. **(a)** PCA of the unit normalized gene expression vectors for the samples results in two clusters divided across PC1. **(b)** Non-negative matrix factorization (NMF) sorted by the coefficients in column 0 with only the first 10 rows shown. Of those only the first three have coefficients larger in column 0 than in column 1.

**Supplementary Figure S2:**
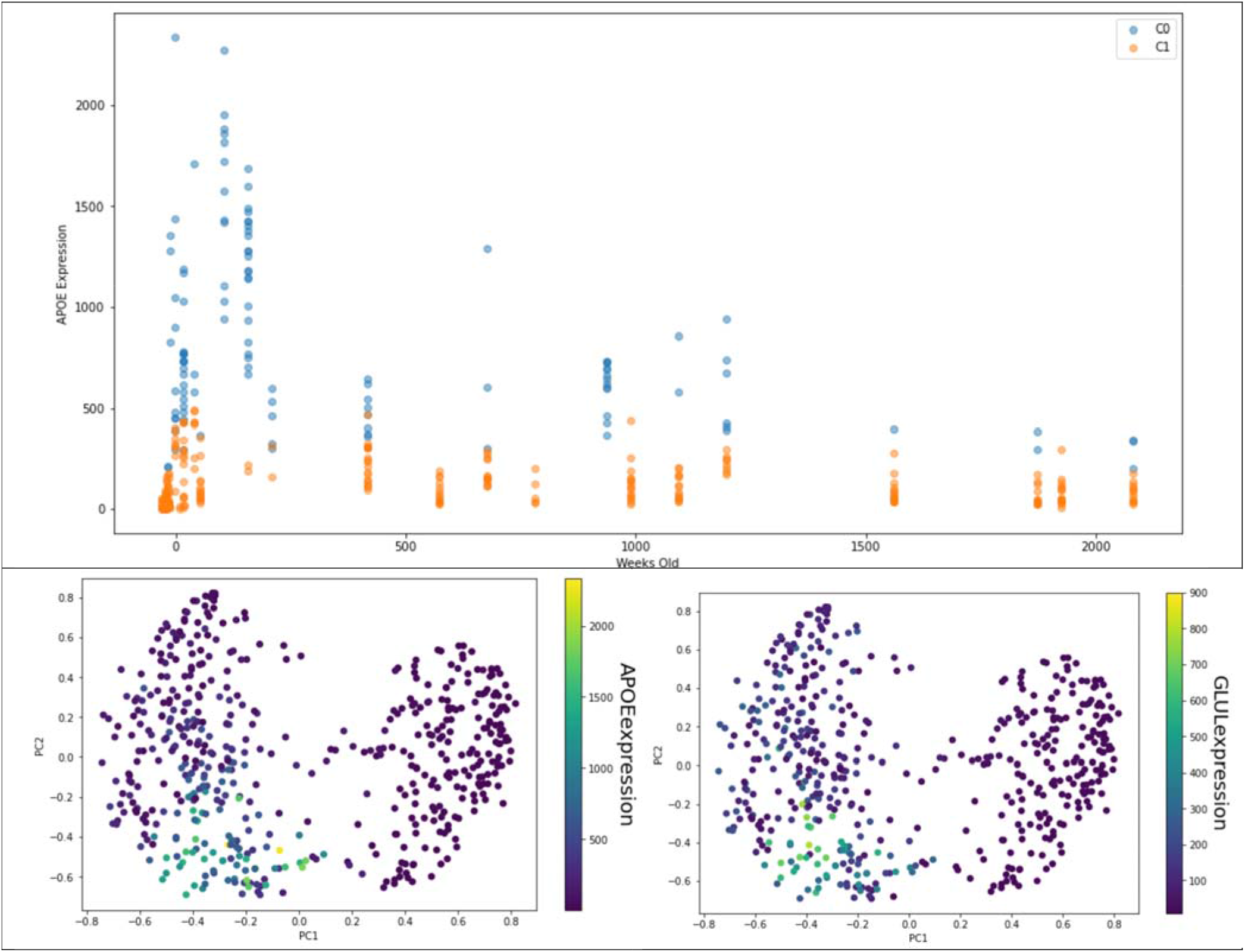
Time series of the two clusters. **(a)** Time series of the two clusters showing the markedly higher expression of APOE in Cluster C0. **(b**,**c)** The strongest APOE and GLUL expression values overlap the same samples as shown by mapping the expression values onto the PCA of Figure S1(a).

**Supplementary Figure S3:**
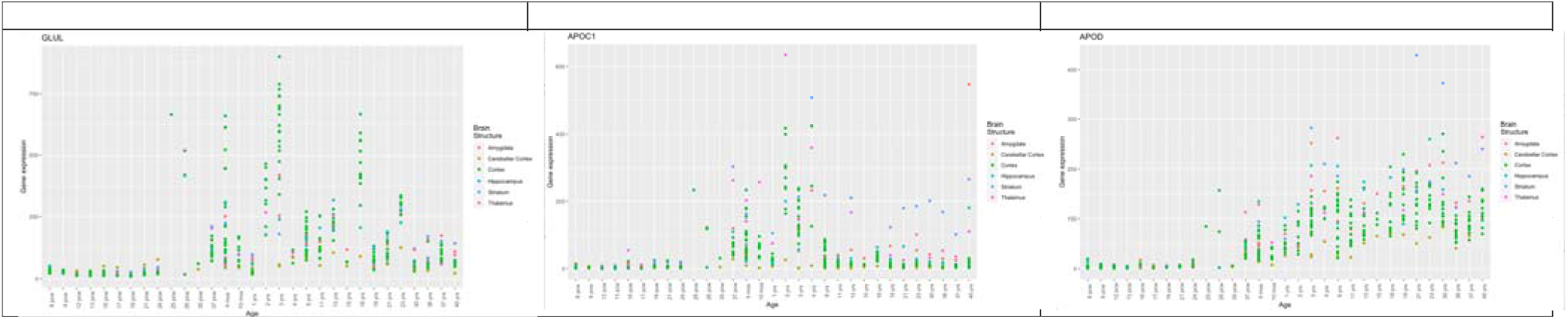
Gene-specific expression pattern over age and brain structures. Note that these scatter plots include all the samples, not just those from one cluster. Each panel corresponds next hightest ranked genes in ***W*** matrix column 0 after APOE (which can be seen in Figure 1). **(a)** GLUL **(b)** APOC1 **(c)** APOD

**Supplementary Figure S4:**
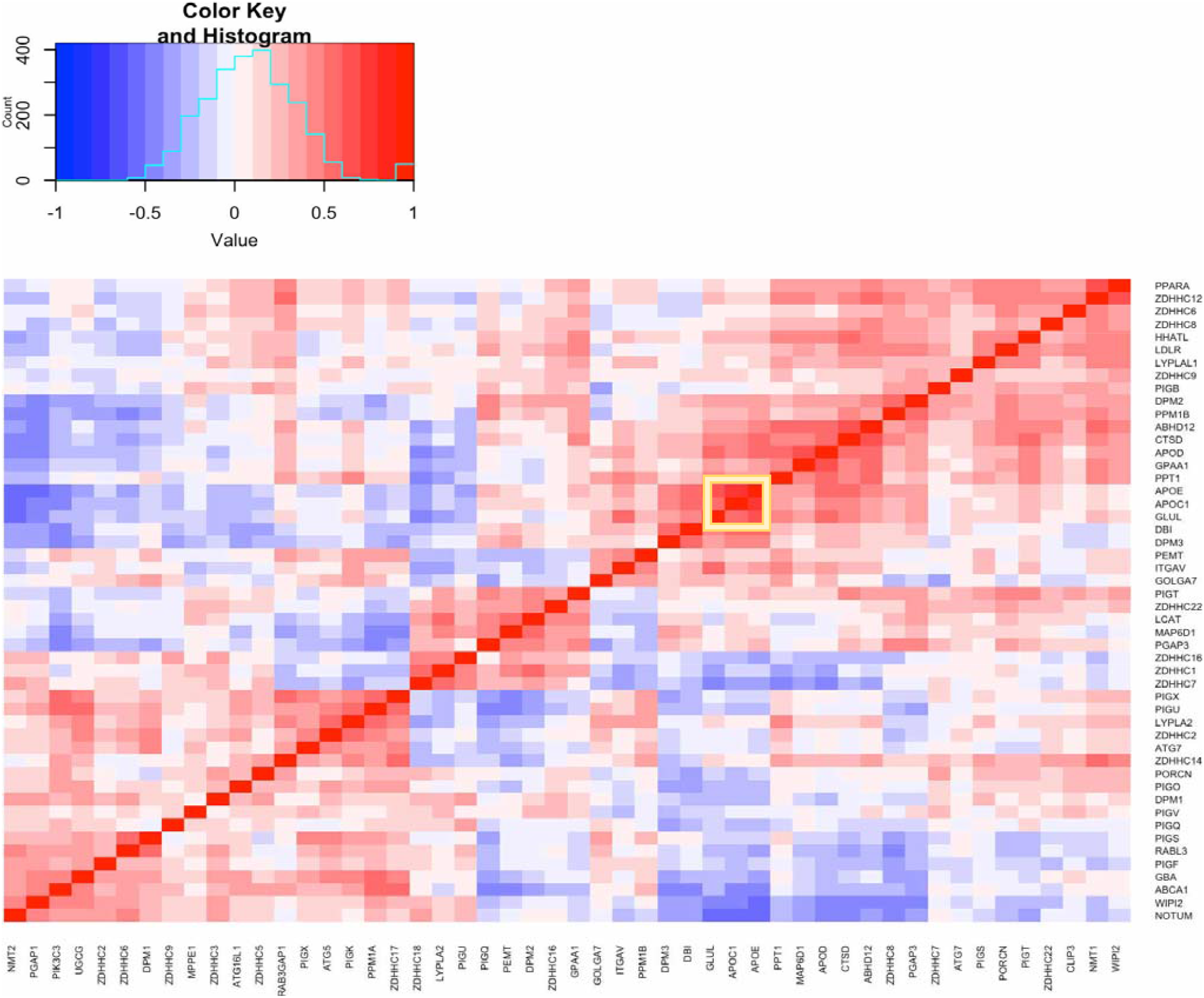
Correlation matrix of lipid gene panel. All genes with median expression < 5 were eliminated to simplify the diagram and all samples were used to calculate the pairwise correlation matrix across genes using the Kendall rank correlation method for robustness. The complete linkage method was for hierarchical clustering and the dendrogram also omitted for simplicity of visualization. As quantified in the Results section, the genes with tightest correlation with APOE are GLUL and APOC1 (correlations outlined by yellow rectangle in the figure).

**Table 1:** Mapping Brainspan structures to a less sparse/more aggregated scheme.

| BRAINSPAN STRUCTURE | AGGREGATED STRUCTURE |
| --- | --- |
| OCX | Cortex |
| M1C-S1C | Cortex |
| AMY | Amygdala |
| MGE | Cortex |
| STC | Cortex |
| URL | Cerebellar Cortex |
| CGE | Cortex |
| DTH | Thalamus |
| MFC | Cortex |
| DFC | Cortex |
| OFC | Cortex |
| LGE | Cortex |
| ITC | Cortex |
| HIP | Hippocampus |
| VFC | Cortex |
| PCX | Cortex |
| TCX | Cortex |
| A1C | Cortex |
| V1C | Cortex |
| STR | Striatum |
| M1C | Cortex |
| IPC | Cortex |
| S1C | Cortex |
| CB | Cerebellar Cortex |
| CBC | Cerebellar Cortex |
| MD | Thalamus |

**Supplementary Table 1.**
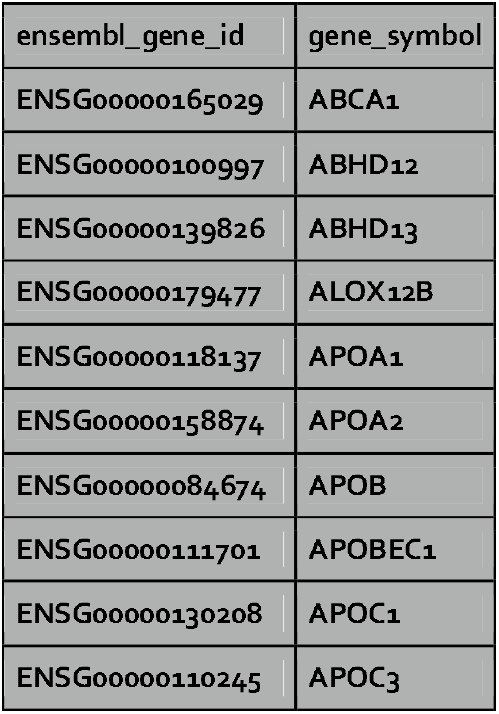
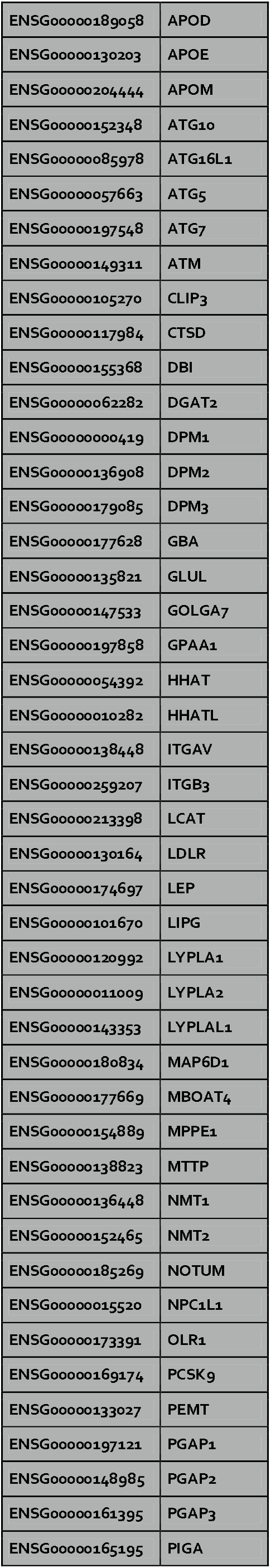
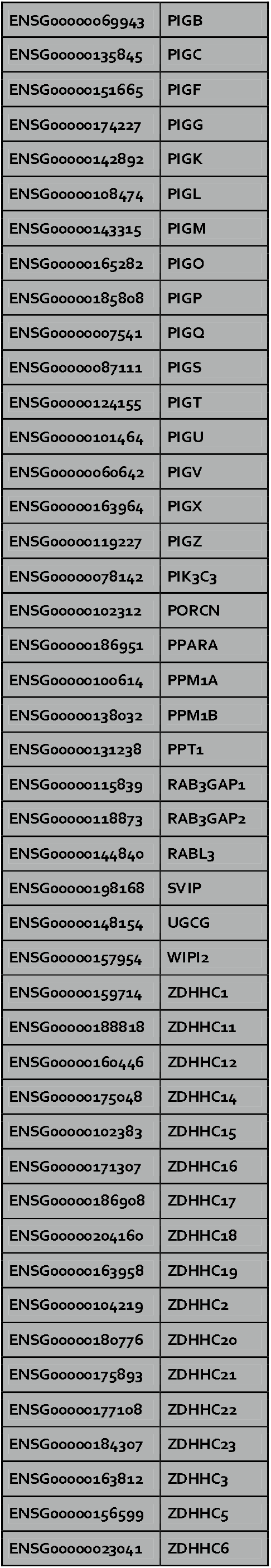
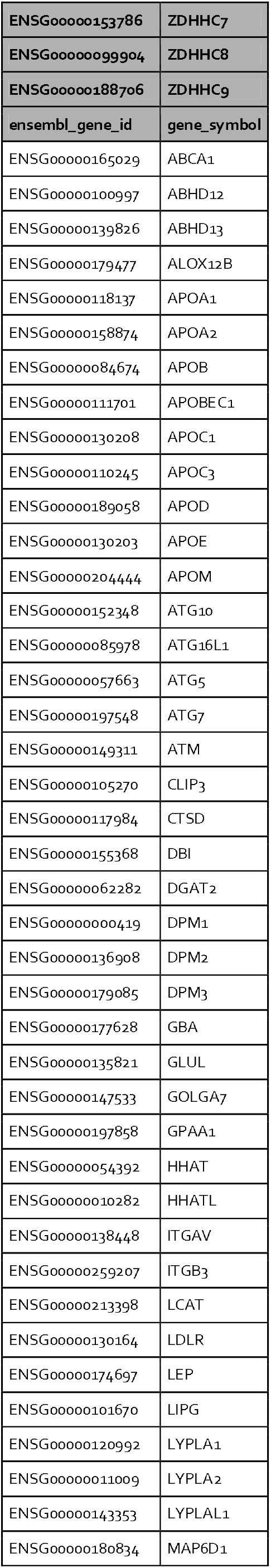
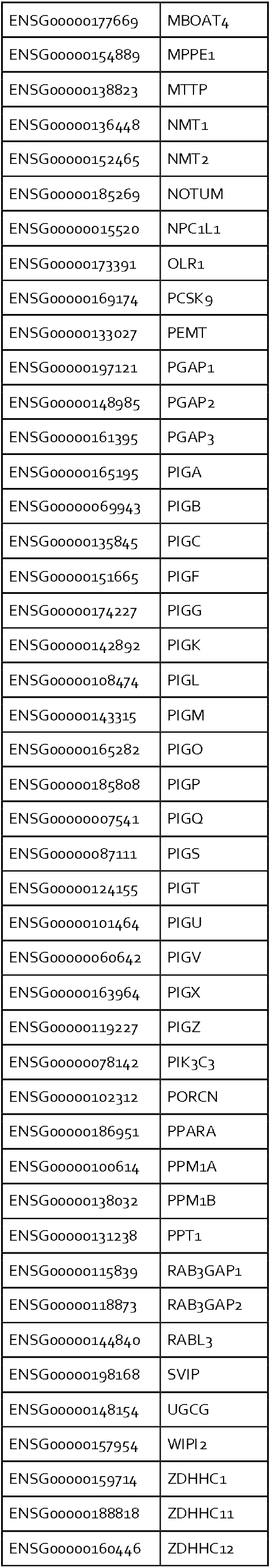
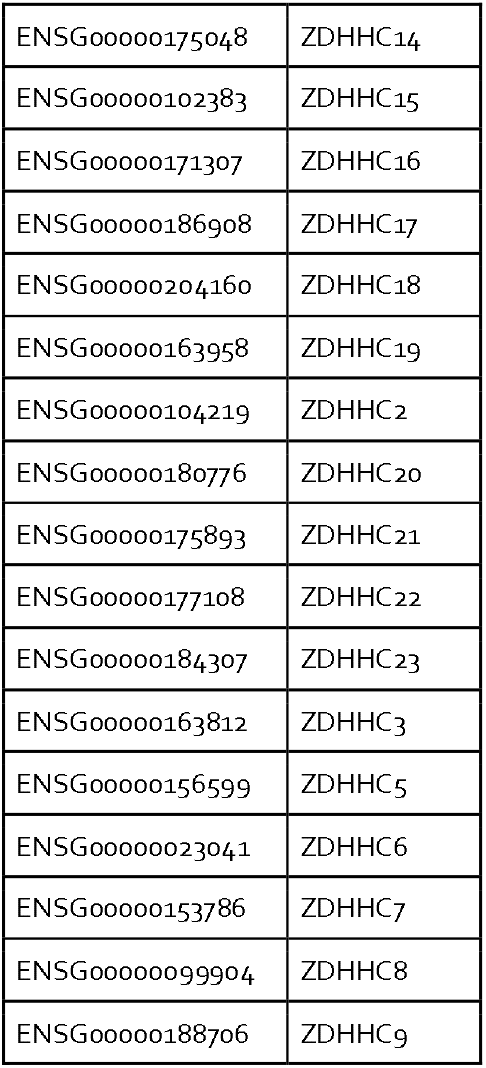

